# Systemic inflammation suppresses lymphoid tissue remodeling and B cell immunity during concomitant local infection

**DOI:** 10.1101/831081

**Authors:** Yannick O Alexandre, Sapna Devi, Simone L Park, Laura K. Mackay, William R. Heath, Scott N. Mueller

## Abstract

Concurrent infection with multiple pathogens occurs frequently in individuals and can result in exacerbated infections and altered immunity. However, the impact of such coinfections on immune responses remains poorly understood. Here we reveal that systemic infection results in an inflammation-induced suppression of local immunity. During localized infection or vaccination in barrier tissues including the skin or respiratory tract, concurrent systemic infection induced a type I interferon-dependent lymphopenia that impairs lymphocyte recruitment to the draining lymph node (dLN). This leads to suppressed lymphoid stromal cell expansion and dLN remodeling and impaired induction of B cell responses and antibody production. Our data suggest that contemporaneous systemic inflammation constrains the induction of regional immunity.

## Introduction

Viral pathogens that infect the skin and mucosa, including herpes simplex virus (HSV) and influenza viruses, induce immune responses in lymph nodes (LN) that drain the site of local infection. Although immune responses to infection with a single pathogen are well defined, people are often infected with multiple pathogens simultaneously and such pathogen coinfections can impair immune responses and alter the outcomes of infections (Mabbott, 2018; Stelekati and Wherry, 2012). Moreover, systemic pathogen infections including HIV, malaria, COVID-19 and bacterial sepsis, as well as other diseases including autoimmune disorders and stroke, result in systemic inflammation that can lead to immunosuppression. Such systemic infections or inflammatory responses are often correlated with reduced protection against localized coinfections (Chang et al., 2013; Edwards et al., 2015; Hotchkiss et al., 2013; Langhorne et al., 2000). Yet, the mechanisms underlying altered immunity during coinfection or inflammation remain poorly understood.

The induction of immune responses relies upon effective recruitment of B cells and T cells from the circulation where they can encounter cognate viral antigens presented by professional antigen presenting cells. Coordinated cellular interactions define T and B cell activation, proliferation and differentiation, and these interactions are supported by specialized microanatomical compartments within LNs (Alexandre and Mueller, 2018). Networks of non-hematopoietic lymphoid stromal cells (LSCs) construct the microarchitecture of LNs, providing both structural and functional support for the induction of immune responses. Lymphatic endothelial cells (LEC) form the afferent and efferent lymphatic vessels required for sampling peripheral tissues and exit of lymphocytes back to the blood circulation, respectively. Lymphocytes that enter the LN from the blood are recruited via specialized blood endothelial cells (BEC) that form PNAd^+^ high endothelial venules (HEV) (Mueller and Germain, 2009). Subsets of mesenchymal fibroblastic reticular cells (FRC) construct the LN architecture and include marginal reticular cells (MRC), follicular dendritic cells (FDC), T cell zone FRC and additional reticular cells localized in B cell follicles. FRC are identified on the basis of expression of podoplanin (PDPN; also called gp38) and lack expression of the endothelial cell marker CD31. FRC respond dynamically to inflammation by modulating their gene expression program that influences immune responses (Gregory et al., 2017; Malhotra et al., 2012). This includes down-regulation of the expression of the chemokines CCL19 and CCL21, which affects the positioning of CD8 and CD4 T cells (Mueller et al., 2007a). The importance of the mesenchymal LSC network was demonstrated in studies showing that depletion of the FRC network results in impaired induction of immune responses (Cremasco et al., 2014; Denton et al., 2014).

Localized infections induce considerable enlargement of the draining LNs to facilitate immune responses, including recruitment and proliferation of lymphocytes and expansion of stromal cell networks. T cell responses are initiated rapidly within LNs, with activation and proliferation of pathogen-specific T cells occurring in as little as 2 days, followed by egress of the expanded pool of effector cells after 5-6 days. B cell responses are initiated with similar rapidity, and B cells differentiate into short-lived antibody secreting cells (ASC) within days, while other B cells migrate to B cell follicles to form germinal centers (GC) and differentiate into high affinity plasma blasts (PB) or memory B cells. GCs form within days after infection and can persist for weeks or months. LN expansion peaks 1-2 weeks after infection, which is delayed relative to the induction of T cell responses, suggesting that remodeling of LN stromal cell networks may be particularly important for supporting B cell responses. We have previously shown that LN remodeling after HSV infection requires recruitment of lymphocytes from the circulation to induce LN expansion, while B cells can sustain stromal cell responses (Gregory et al., 2017).

Here, we examined the impact of systemic infection on an ongoing local immune response. We show that the induction of systemic inflammation in response to coinfection of mice with lymphocytic choriomeningitis virus (LCMV) or Toll-like receptor (TLR) agonists impaired LN swelling and stromal cell remodeling and inhibited B cell responses to localized infection or vaccination. Systemic inflammation induced a type I interferon (IFN-I) dependent lymphopenia, impairing the recruitment of B cells to draining LNs and hindering the induction of humoral immune responses. Our findings reveal that a consequence of an acute systemic inflammatory response is the suppression of the ability to induce local immunity.

## Results

### Restrained LN remodeling and suppressed B cell responses after LCMV infection

We first investigated how different viral pathogens influence lymphoid tissue remodeling. We infected mice subcutaneously in the footpad with HSV or LCMV Armstrong and enumerated stromal cell populations in the draining popliteal LN (pLN) by flow cytometry 8 days after infection (Fig. 1A). HSV inoculation results in an infection that remains localized to tissues near the site of inoculation (Gregory et al., 2017), whereas LCMV replicates locally as well as spreading systemically, including to the spleen (Jones et al., 2000; Olson et al., 2012). HSV infection induced a marked increase in total pLN cellularity and expanded all populations of pLN LSC: PDPN^+^ FRC, PDPN^+^ CD31^+^ LEC, CD31^+^ BEC and CD31^+^ PNAd^+^ HEV (Fig. 1B). In contrast, LCMV infection resulted in only a marginal increase in cellularity and no significant expansion of the subsets of LSC in the LN compared to uninfected mice (Fig. 1B).

**Fig. 1.**
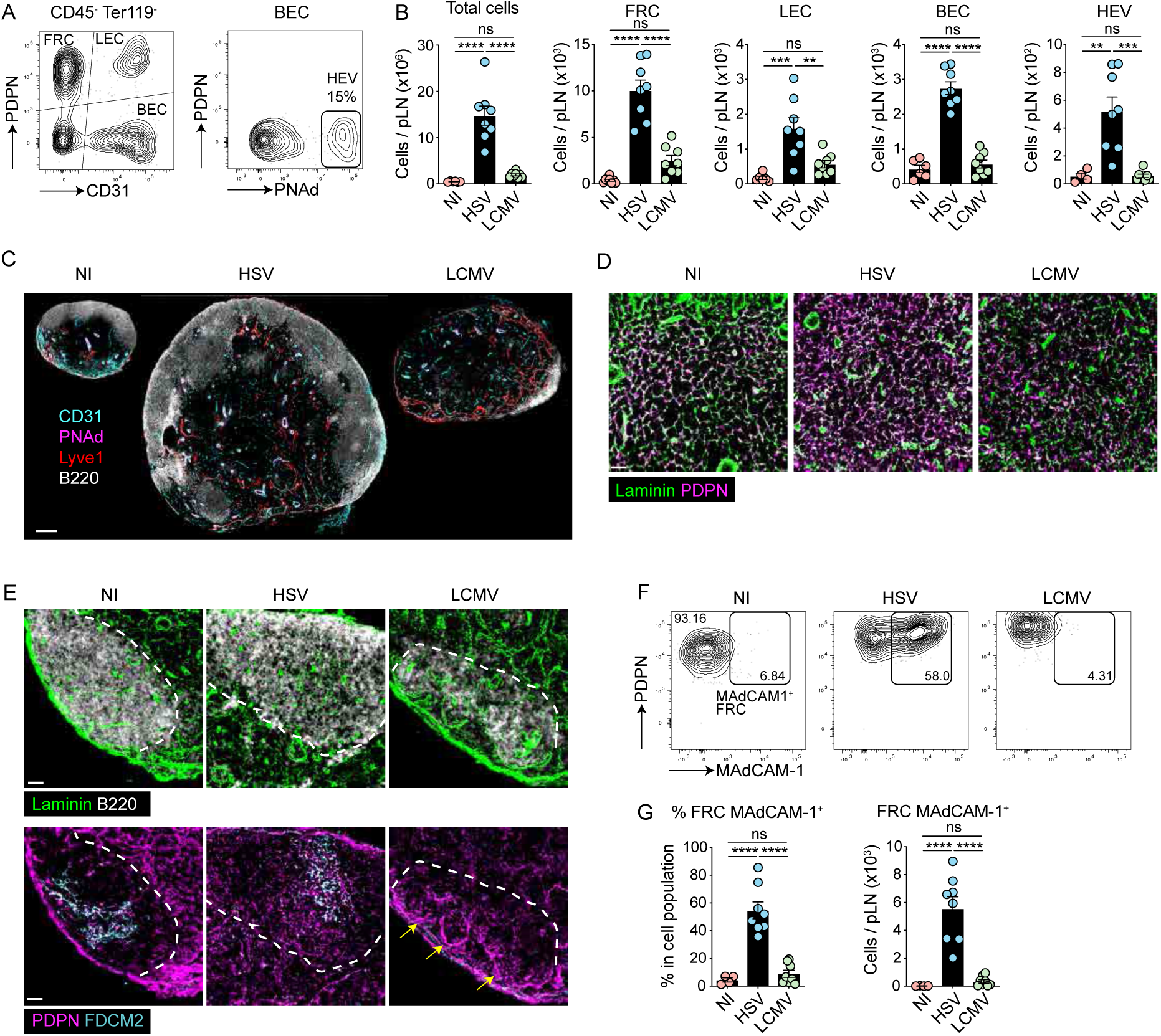
Restrained remodeling of the draining LN during LCMV infection. (A) Gating strategy to identify CD45^−^ Ter119^−^ stromal cell subsets by flow cytometry. (B) Numbers of total cells and stromal cell subsets in the pLN from uninfected (NI) mice and mice infected s.c. with HSV or LCMV analyzed by flow cytometry. Graphs show pooled data (mean ± SEM) from 3 independent experiments each with 2-3 mice per group. *P < 0.05, **P < 0.01, *** P < 0.001, ****P < 0.0001, ns, non-significant, by ANOVA with Tukey’s multiple comparisons test. (C-E) pLN sections from NI, HSV or LCMV infected mice were stained for B220, Lyve1, CD31 and PNAd (C), laminin and podoplanin (D) or laminin, B220, podoplanin, amd FDC-M2 (E) and analyzed by confocal microscopy. Data are representative of 2 experiments with 3 mice per group. Dotted lines depict border of B cell follicles and arrows show subcapsular FRC in LCMV pLN that were not observed in NI or HSV pLN. Scale bars, 200μm (C), 30μm (D-E) (F) MAdCAM-1 expression on FRC from NI mice and following HSV or LCMV infection. (G) Percentage and numbers of MAdCAM-1^+^ FRC in the pLN of NI and mice infected s.c. with HSV or LCMV. Graphs show pooled data (mean ± SEM) from 3 independent experiments each with 2-3 mice per group. ****P < 0.0001, ns, non-significant, by ANOVA with Tukey’s multiple comparisons test.

Examination of pLN by confocal microscopy revealed substantial enlargement of the draining LN after HSV infection, whereas pLN were small and lacked organized B cell zones after LCMV infection (Fig. 1C). Closer examination revealed substantial disorganization of the PDPN^+^ T cell zone FRC network that ensheaths laminin^+^ reticular fibers (Fig. 1D) and loss of B cell zone stromal cell organization after LCMV infection (Fig. 1E). Notably, in LCMV infected mice B cell follicles were highly disorganized, and we could not detect FDC, whereas B cell follicle organization and FDCs were intact in HSV infected mice (Fig. 1E). Moreover, we observed the formation of a network of PDPN^+^ FRC on the capsular side of the B cell follicles in LCMV infected mice (Fig. 1E, yellow arrows). This indicated that the B cell follicles retracted from the LN capsule and a population of subcapsular PDPN^+^ FRC formed in this zone during LCMV infection.

Because MAdCAM-1^+^ MRC form a specialized stromal layer between B cell follicles and the subcapsular sinus, we postulated that the expanded subcapsular LSC in LCMV infected mice might be MRC, but subsequent analysis revealed that MAdCAM-1^+^ were not expanded following LCMV infection. Conversely, we observed a substantial increase in MAdCAM-1^+^ PDPN^+^ FRC after HSV infection (Fig. 1F). Greater than half of all FRC were MAdCAM-1^+^ after HSV infection (Fig. 1G), suggesting that expression of this adhesion molecule was upregulated after HSV but not LCMV infection. Staining of tissue sections confirmed increased expression of MAdCAM-1 on T cell zone FRC in pLN after HSV but not LCMV infection (Fig. S1). This suggests that MAdCAM-1, which is typically used to define MRC and is also expressed by FDC in LNs, is not a reliable marker of MRC after infection.

Next, we examined virus-specific CD8^+^ T cell responses in the pLN by major histocompatibility complex class I (MHC-I) tetramer staining. Local infection with LCMV induced larger numbers of virus-specific CD8^+^ T cells compared to HSV infection (Fig. 2A-B). In contrast, we found that B cell numbers were significantly reduced in draining pLN after LCMV infection in comparison with HSV infected mice (Fig. 2D). Expansion of GL7^+^ germinal center (GC) B cells and CD138^+^ antibody secreting cells (ASC) was also significantly lower in the pLN after LCMV infection compared to HSV infection (Fig. 2C-E). B cell follicles were increased in size after HSV infection and contained GL7^+^ GC B cells, while B cell follicles were markedly reduced in size and GL7^+^ cells were extrafollicular in LCMV infected mice (Fig. 2F). Large numbers of CD138^+^ ASC accumulated at the border between T and B cell zones and within the medullary cords after HSV infection, whereas ASC were reduced and only observed in the medulla during LCMV infection (Fig. 2F). Thus, LCMV infection results in diminished LN remodeling, loss of B cell follicles and restrained B cell responses in draining LN.

**Fig. 2.**
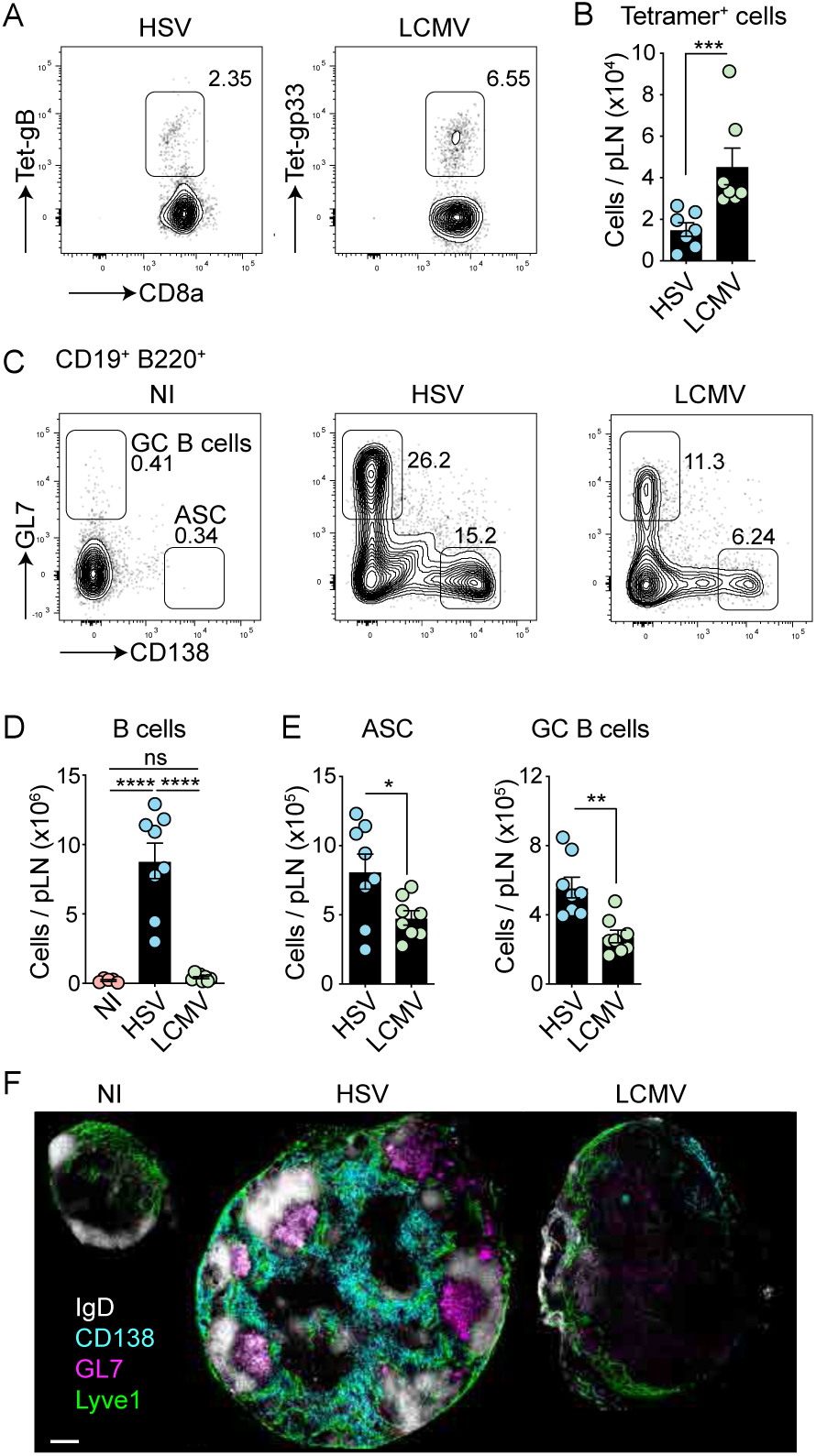
Suppressed B cell responses during LCMV infection. (A) Tetramer staining of CD8^+^ T cells, following HSV and LCMV infection. (B) Numbers of CD8^+^ Tetramer^+^ cells in the pLN of HSV and LCMV-infected mice. Data are pooled from 2 independent experiments with 3-4 mice per group. ***P < 0.001, by Mann-Whitney test. (C) Flow cytometry analysis of GL7^+^ GC B cells and CD138^+^ ASC. B cells were first gated on live CD19^+^ CD3^−^ NK1.1^−^ cells. (D) Absolute numbers of B cells in the popliteal LN from NI mice and mice infected with HSV or LCMV. Graphs show pooled data (mean ± SEM) from 3 independent experiments each with 2-3 mice per group. ****P < 0.0001, ns, non-significant, by ANOVA with Tukey’s multiple comparisons test. (E) ASC and GC B cells in the pLN from mice infected with HSV or LCMV. Graphs show pooled data (mean ± SEM) from 2 independent experiments each with 4 mice per group. *P < 0.05, **P < 0.01, by unpaired two-tailed t test. (F) pLN sections from NI, HSV or LCMV infected mice were stained for IgD, CD138, GL7 and Lyve1 and analyzed by confocal microscopy. Data are representative of 2 experiments with 3 mice per group. Scale bar, 200μm.

### LCMV coinfection suppresses LN remodeling in response to localized infection

The subdued stromal cell expansion and altered lymphoid tissue architecture observed following LCMV infection led us to investigate the impact of LCMV infection on LN remodeling during localized HSV coinfection. We infected mice with HSV in the footpad and coinfected the mice 2 days later with LCMV either by the same route or systemically by intraperitoneal infection (Fig. 3A). LCMV coinfection by either route markedly restrained the increase in pLN cellularity induced by HSV infection and suppressed FRC expansion (Fig. 3B). Local infection with LCMV resulted in infection of pLN FRC whereas systemic LCMV infection did not (Fig. S2A). Yet, both routes of infection resulted in equivalent suppression of the draining LN response (Fig. 3B), ruling out an inhibitory role of LCMV through direct infection of FRC. Since local LCMV infection also results in systemic spread of virus from the LN to spleen (Olson et al., 2012), we performed subsequent experiments using systemic LCMV coinfection. Importantly, LCMV coinfection did not alter the growth of HSV in the tissue (Fig. 3C).

**Fig. 3.**
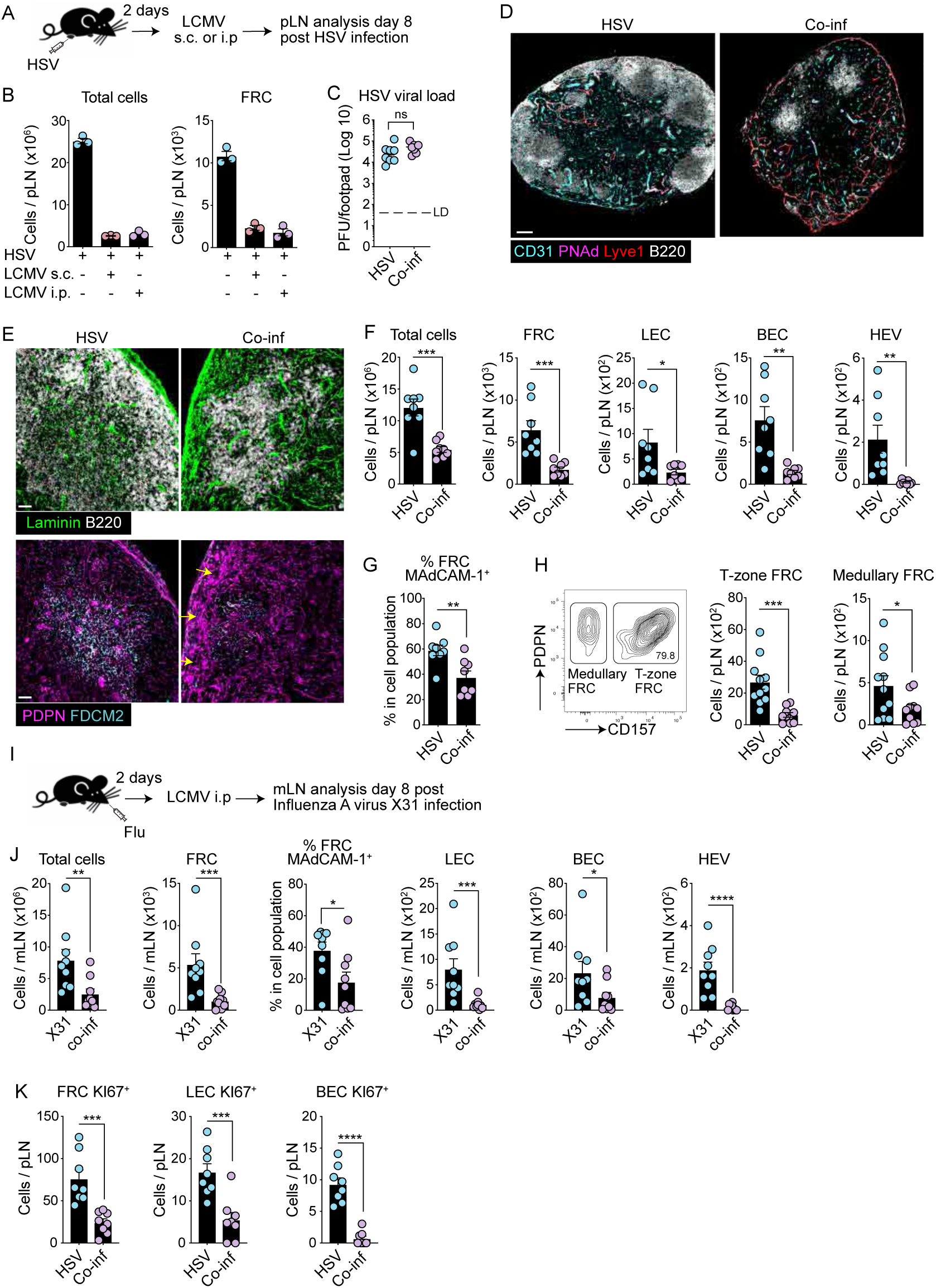
LCMV coinfection suppresses lymphadenopathy to localized infection. (A) Experimental schematic of coinfection. Mice were infected subcutaneously (s.c.) with HSV and 2 d later infected with LCMV s.c. or intraperitoneally (i.p.). pLN were analyzed 8 d post HSV infection. (B) Numbers of total cells and FRC in the pLN of infected mice analyzed by flow cytometry. Graphs are representative of 2 experiments with 2-3 mice per group (mean ± SEM). (C) HSV viral load in the footpad of infected mice. Mice were infected as in A and footpad tissue harvested 5 d post-HSV infection for plaque assay. Data are pooled from 2 independent experiments with 4 mice per group. ns, non-significant, by Mann-Whitney test LD: limit of detection. (D-E) pLN sections from NI, HSV or coinfected mice were stained for B220, Lyve1, CD31, and PNAd (D), Laminin, B220, podoplanin, and FDC-M2 (E) and analyzed by confocal microscopy. Data are representative of 2 experiments with 3 mice per group. Scale bar, 40μm. (F-G) Numbers of total cells and stromal cell subsets (F) and MAdCAM-1 expression analysis on FRC (G) in the pLN of HSV and coinfected mice analyzed by flow cytometry. Graphs show pooled data (mean ± SEM) from 2 independent experiments with 4 mice per group. (H) Expression of CD157 on PDPN^+^ CD31^−^ MAdCAM-1^−^ CD21/35^−^ cells to define T cell zone FRC (CD157^+^) and medullary FRC (CD157^−^) by flow cytometry. Graphs are pooled data of 3 experiments with 2-4 mice per group (mean ± SEM). *P < 0.05, **P < 0.01, *** P < 0.001, ****P < 0.0001, by unpaired two-tailed t test (F-H). (I) Mice were infected intranasally with Flu-gB and 2 d later infected systemically with LCMV. Mediastinal LN (medLN) were analyzed 8 d post Flu infection. (J) Numbers of total cells, stromal cell subsets, and MAdCAM-1 expression analysis on FRC in the medLN of Flu and coinfected mice analyzed by flow cytometry. Graphs show pooled data (mean ± SEM) from 2 independent experiments with 4-5 mice per group. *P < 0.05, **P < 0.01, *** P < 0.001, ****P < 0.0001, by Mann-Whitney test. (K) Ki67^+^ stromal cell subsets following HSV and coinfection. Mice were infected as in A and pLN harvested 5 d post-HSV infection and intracellular KI67 expression was analyzed flow cytometry. Graphs show pooled data (mean ± SEM) from 2 independent experiments each with 4 mice per group. *** P < 0.001, ****P < 0.0001, by unpaired two-tailed t test.

We examined HSV-draining pLN sections by confocal microscopy and discovered that LCMV coinfection prominently altered pLN remodeling in response to HSV. In coinfected mice, B cell follicles were small and disorganized and LYVE-1^+^ medullary sinuses were more prominent compared to mice infected with HSV alone (Fig. 3D). Closer examination of B cell follicles in coinfected mice revealed fewer B cells, reduction of the FDC network within the follicles and recession of the follicles away from the pLN subcapsular sinus (Fig. 3E). Strikingly, a dense network of PDPN^+^ FRC formed between B cell follicles and the pLN capsule in coinfected mice (Fig. 3E, yellow arrows), similar to what we observed during local LCMV infection (Fig. 1E), that was absent from mice infected with HSV alone.

Quantification of cells by flow cytometry revealed a significant reduction in total cellularity and reduced expansion of pLN FRC, LEC, BEC and PNAd^+^ HEV in HSV-draining LN from LCMV coinfected mice (Fig. 3F). Coinfection suppressed the induction of MAdCAM-1 on FRC (Fig. 3G), resulting in a significant reduction in both MAdCAM-1^+^ as well as MAdCAM-1^−^ FRC (Fig. S2B). We also examined CD157 (BP-3) expression on FRC, a marker that has been used to identify subpopulations of FRC including CD157-lo medullary FRC (Huang et al., 2018). Both CD157^+^ T cell zone FRC and CD157-lo medullary FRC populations were reduced in the pLN of mice coinfected with HSV and LCMV in comparison with mice infected with HSV alone (Fig. 3H).

These data suggested that LCMV coinfection inhibited LN expansion in response to local HSV infection. To determine if systemic coinfection also suppressed dLN responses to other localized infections, we infected mice intranasally with Influenza A virus X31 and coinfected these mice with LCMV 2 days later (Fig. 3I). Coinfection suppressed immune responses in the draining mediastinal LNs (mLN), resulting in reduced cellularity and impaired expansion of FRC, LEC, BEC and PNAd^+^ HEV, as well as impaired upregulation of MAdCAM-1 by mLN FRC compared to mice infected with influenza alone (Fig. 3J). This indicated that systemic infection concurrent with either a skin or respiratory viral infection suppresses immune responses in the LN draining the site of localized challenge.

Since systemic coinfection suppressed expansion of pLN LSC, we wanted to know if this was due to reduced proliferation or altered survival of LSC. We examined proliferating LSC in LCMV and HSV coinfected mice by staining for Ki67. Significantly fewer Ki67^+^ FRC, LEC and BEC were detected in coinfected mice when compared to mice infected with HSV alone (Fig. 3K). In contrast, coinfection did not alter the number of apoptotic (Annexin-V+ PI+) FRC, LEC or BEC compared to that observed following infection with HSV alone (Fig. S2C). Further, we observed no alterations in metabolic functions as determined by glucose or lipid uptake, mitochondrial mass or oxidative stress in pLN LSC in coinfected mice compared to mice infected with HSV alone (Fig. S2D-E). Thus, systemic LCMV coinfection suppressed dLN remodeling and stromal cell expansion in response to localized infection.

### Systemic coinfection suppresses B cell responses to localized infection

We examined immune responses in HSV and LCMV coinfected mice and observed that although total numbers of CD8^+^ T cells in the pLN were reduced by coinfection, HSV-specific CD8^+^ T cell responses were unaffected (Fig. 4A). Systemic LCMV coinfection also reduced the accumulation of total CD4^+^ T cells in the draining pLN and suppressed the induction of T follicular helper (T_FH_) cells compared to mice infected with HSV alone (Fig. 4B, S3A). B cell numbers were significantly reduced in the pLN of coinfected mice and the induction of GC B cells and ASCs was impaired (Fig. 4C). This suppression of local B cell responses during LCMV coinfection was not due to increased apoptosis of B cells or T cells, as indicated by AnnexinV staining (Fig. 4D). Histological examination of pLN sections revealed a reduction in the size and numbers of GL7^+^ GC B cells and fewer CD138^+^ plasma cells in coinfected mice (Fig. 4E). Moreover, systemic LCMV coinfection significantly suppressed HSV-specific antibody responses (Fig. 4F).

**Fig. 4.**
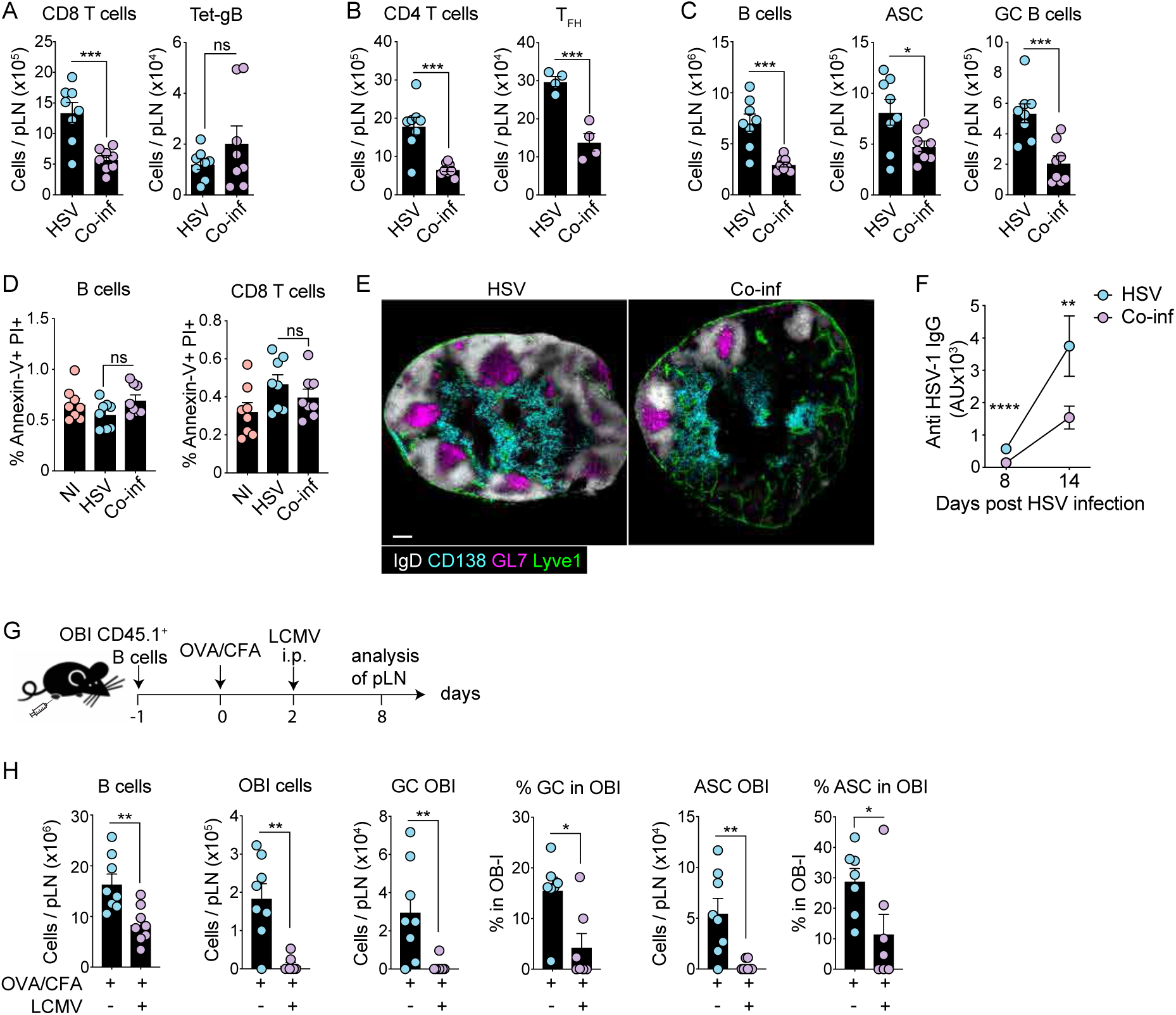
Systemic coinfection suppresses B cell responses to localized infection. (A-C) Mice were infected as in Fig. 3A and pLN analyzed for CD8^+^ T cells and HSV gB tetramer^+^ cells (A), CD4^+^ T cells and CXCR5^+^ PD1^+^ CD4^+^ T_FH_ cells (B) and B cells (C). Graphs show pooled data (mean ± SEM) from 1-2 independent experiments with 4 mice per group. *P < 0.05, *** P < 0.001, ****P < 0.0001, ns, non-significant, by unpaired two-tailed t test. (D) Coinfection does not increase lymphocyte cell death. pLN from HSV and coinfected mice were harvested 5 d post-infection and analyzed for Annexin V and PI expression on B and CD8^+^ T cells by flow cytometry. Inguinal LN from naïve mice were included as controls. Graphs show pooled data (mean ± SEM) from 2 independent experiments with 4 mice per group. ns, non-significant, by ns, non-significant, by unpaired two-tailed t test (E) pLN sections from NI, HSV or LCMV infected mice were stained for IgD, CD138, GL7 and Lyve1 and analyzed by confocal microscopy. Data are representative of 2 experiments with 3 mice per group. Scale bar, 200μm. (F) Serum anti–HSV-1 IgG titer assessed by ELISA. Mice were infected as in Fig. 3 and sera harvested at day 8 and 14 post-HSV infection. Graph shows pooled data (mean ± SEM) from 2 independent experiments with 4 mice per group. *P < 0.05, ** P < 0.01, by unpaired two-tailed t test. (G) Experimental schematic. Mice were given CD45.1^+^ OBI B cells 1 day prior to footpad immunization with OVA/CFA. Mice were infected with LCMV i.p. 2 days later. pLN were analyzed 8 d. (H) OBI B cell response in the pLN. Mice were injected i.v. with transgenic OBI-GFP cells and two days later injected s.c. with an emulsion of OVA/CFA. Mice were infected i.p. with LCMV 2 d post immunization and pLN harvested 8 d post OVA/CFA injection. Graphs show pooled data (mean ± SEM) from 2 independent experiments with 4 mice per group. *P < 0.05, ** P < 0.01, by Mann-Whitney test.

To further investigate the impact of systemic coinfection on the induction of local antigen-specific B cell responses, we examined ovalbumin (OVA)-specific B cell receptor transgenic OB-I B cell responses following transfer into B6 mice and immunization with OVA in complete freunds adjuvant (OVA/CFA) (Fig, 4G). OB-I B cells expanded significantly in the pLN following sub-cutaneous OVA/CFA immunization and formed GC B cells and ASCs (Fig. 4H and Fig S3B). Coinfection of mice with LCMV significantly impaired the expansion of the OB-I B cells in the pLN, reduced the number of GC cells and ASCs and reduced the frequency of OB-I B cells that developed into GC cells and ASCs (Fig. 4H and Fig. S3B). Together, these data show that systemic coinfection with an unrelated pathogen suppresses the induction of antigen-specific B cell responses in LN draining the site of a localized challenge.

### Coinfection-induced lymphopenia suppresses LN expansion and B cell responses

CTL can kill virus infected stromal cells and B cells during infection of mice with strains of LCMV that cause chronic infection (Moseman et al., 2016; Mueller et al., 2007b). To determine if CD8^+^ T cells contributed to reduced LN expansion and B cell responses during coinfection with LCMV Armstrong, which is an acute infection, we infected mice with HSV then treated with depleting anti-CD8α antibody prior to LCMV coinfection (Fig. 5A). Suppression of pLN remodeling and impairment of B cell responses in LCMV coinfected mice was unaffected by the loss of CD8^+^ T cells (Fig. 5B and Fig. S3C). Inflammatory monocytes can impair B cell responses during chronic LCMV infection (Sammicheli et al., 2016), but B cell responses remained constrained with LCMV coinfection in CCR2^−/−^ hosts with defective recruitment of monocytes (Fig. S3D).

**Figure 5.**
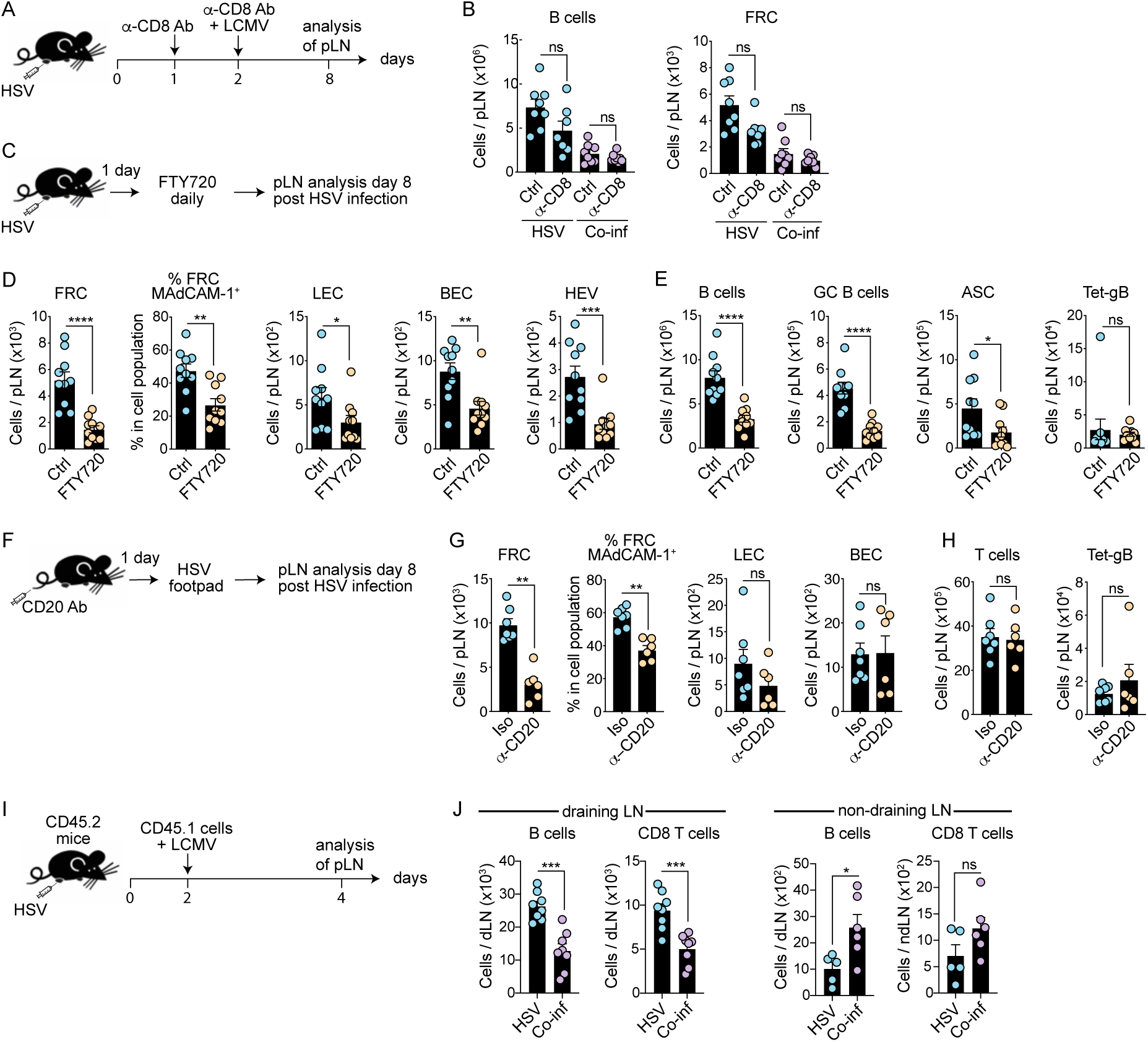
Lymphopenia induced during systemic infection suppresses local LN responses. (A) Experimental schematic of CD8^+^ T cell depletion during coinfection. Mice were infected s.c. with HSV and injected with CD8 depleting antibody on days 1 and 2. Mice were infected with LCMV on day 2. pLN were analyzed 8 d post HSV infection. (B) Numbers of B cells and FRC in the pLN of mice infected with HSV or coinfected with or without CD8^+^ T cells. Graphs show pooled data (mean ± SEM) from 2 independent experiments with 3-4 mice per group. ns, non-significant, by ANOVA with Krukal-Wallis test. (C) Experimental schematic of FTY720 treatment during HSV infection. Mice were infected s.c. with HSV and the following day injected i.p. with FTY720 or control for 7 d. pLN were analyzed 8 d post-HSV infection. (D-E) Absolute numbers of stromal cell subsets and MAdCAM-1 expression on FRC (D), B and T cell responses (E) in pLN of HSV infected mice treated with FTY720. Graphs show pooled data (mean ± SEM) from 2 independent experiments with 5 mice per group. (F) Experimental schematic of B cell depletion during HSV infection. Mice were injected i.p. with CD20 depleting antibody and the following day infected s.c. with HSV. (G-H) Analysis of stromal cells and T cells by flow cytometry in HSV-infected B cell-depleted mice. Graphs show pooled data (mean ± SEM) from 2 independent experiments with 3-4 mice per group. (I-J) Adoptive transfer of congenic lymphocytes into HSV infected mice on day 2 and LCMV coinfection, with analysis of CD45.1^+^ cell numbers in the draining popliteal LN and non-draining LN 2 days later. *P < 0.05, **P < 0.01, *** P < 0.001, ****P < 0.0001, ns, non-significant, by Mann-Whitney test (D-E, G-H, J).

The reduction in B and T lymphocytes in the pLN of LCMV coinfected mice led us to hypothesize that LCMV infection induces a general lymphopenia that traps lymphocytes in distal tissues and inhibits lymphocyte recruitment to the draining LN during peripheral viral infection. To examine this, we treated HSV-infected mice with the immunomodulatory drug FTY720 to induce a systemic lymphopenia (Halin et al., 2005) (Fig. 5C). Expansion of pLN LSC, and upregulation of MAdCAM-1 by FRC was impaired in FTY720 treated HSV infected mice compared to untreated controls (Fig. 5D). B cell accumulation and induction of GC B cells and ASC was also inhibited in FTY720 treated mice, whereas virus-specific CD8^+^ T cell responses were unchanged, suggesting that lymphopenia impairs the induction of local B cell responses and LN remodeling but not virus-specific CD8^+^ T cell responses (Fig. 5E).

To determine if B cell recruitment was required for expansion of the LN, we treated mice with an anti-CD20 antibody to deplete B cells prior to HSV infection and examined stromal cell and T cell responses in B cell ablated mice (Fig. 5F and Fig. S3E). In the absence of B cells, FRC expansion was significantly reduced and upregulation of MAdCAM-1 by FRC was blunted, whereas LEC and BEC expansion was unaffected (Fig. 5G). T cell recruitment to the HSV-draining pLN and induction of HSV-specific CD8^+^ T cell responses were unimpaired by depletion of B cells (Fig. 5H). Thus, recruitment of B cells to the dLN during localized viral infection is required to drive FRC expansion, while additional signals contribute to the remodeling of LN endothelial cell compartments.

To examine if systemic coinfection impaired cell recruitment to the dLN, we adoptively transferred naïve lymphocytes into HSV and LCMV coinfected mice (Fig. 5I). B cell and T cell recruitment to the draining LN was suppressed, with a concomitant increase in cell recruitment to distal LNs (Fig. 5J). Together, these data show that systemic coinfection impairs lymphocyte recruitment to regional dLN and impairs local LN remodeling.

### Systemic inflammation suppresses local immunity via IFN-I

We hypothesized that systemic inflammation was responsible for the lymphopenia-induced suppression of local LN responses. To examine this, we administered the TLR agonists poly(I:C) or lipopolysaccharide (LPS) to HSV infected mice (Fig. 6A). Both inflammatory mediators substantially reduced the HSV-driven increase in pLN cellularity, FRC expansion, as well as B cell responses (Fig. 6B). Systemic LCMV infection induces the production of pro-inflammatory cytokines, including type I interferon (IFN-I) (Doughty et al., 2001). LCMV induced a substantial lymphopenia in mice that was dependent on IFN-I receptor (IFNAR) signaling as it did not occur in *Ifnar2*^−/−^ mice (Fig. S4A). We asked whether IFNAR signals contributed to reduced LN hypertrophy and B cell recruitment during coinfection. We generated chimeric mice with a mixture of B cell deficient (μMT^−/−^) and *Ifnar2*^−/−^ BM to confine the lack of IFNAR expression to B cells (Fig. S4B). However, loss of IFNAR in B cells did not restore LN remodeling or B cell responses in coinfected mice (Fig. S4C). Examination of reciprocal BM chimeric mice generated by reconstitution of either WT or *Ifnar2*^−/−^ mice with WT or *Ifnar2*^−/−^ BM revealed that cells in both the hematopoietic and non-hematopoietic compartments contributed to altered local LN responses via IFNAR signaling (Fig. 6C and S4D-E).

**Figure 6.**
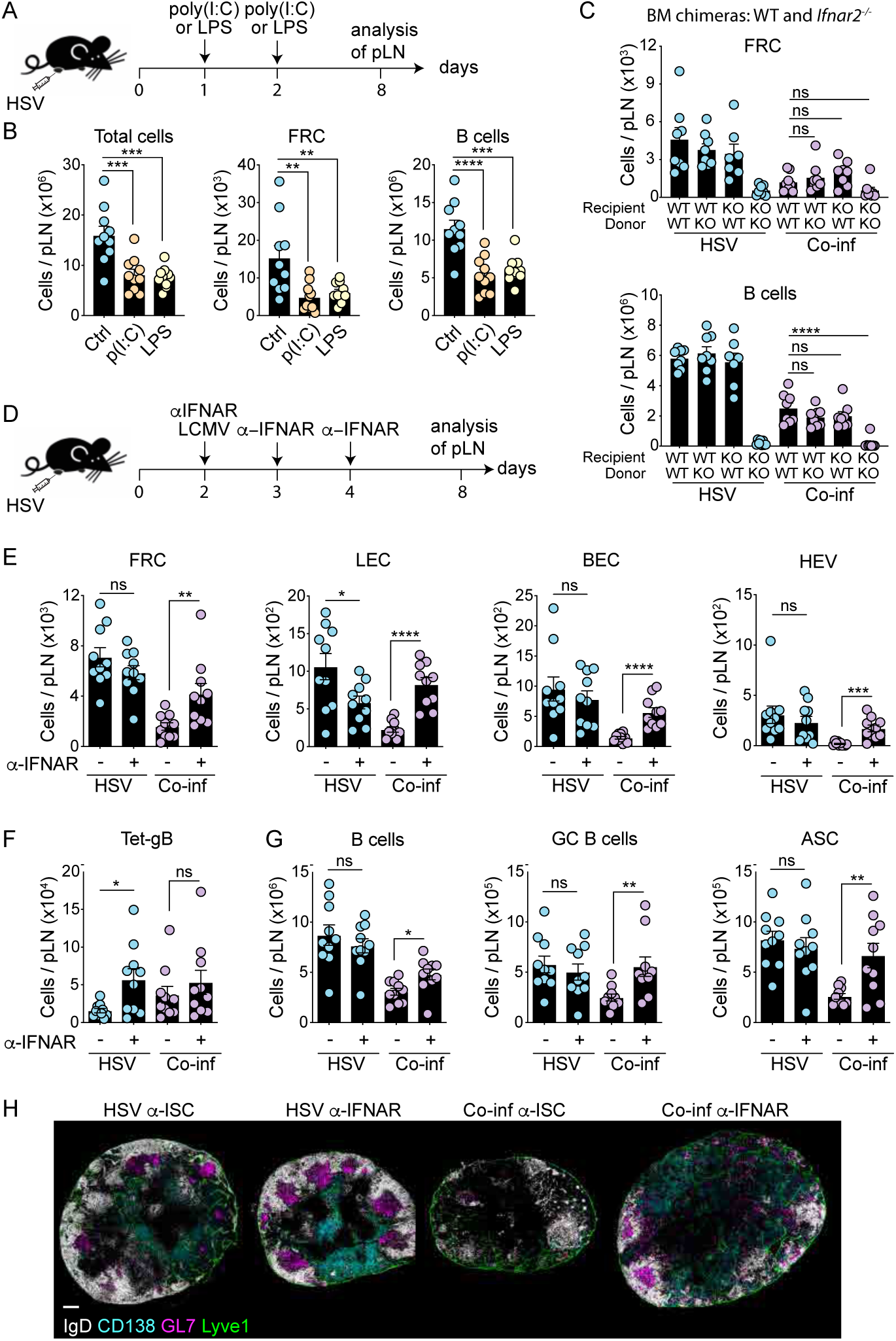
Systemic inflammation suppresses local immunity via IFN-I. (A) Experimental schematic of adjuvant injection in HSV-infected mice. Mice were infected s.c. with HSV and 2 d later received 2 i.p. injections of either poly(I:C) or LPS 24 h apart. pLN were harvested 8 d post-HSV infection. (B) Numbers of total cells, FRC and B cells analyzed by flow cytometry in infected mice injected with poly(I:C) or LPS. Graphs show pooled data (mean ± SEM) from 2 independent experiments with 5 mice per group. (C) Cell numbers in the pLN of HSV and coinfected WT and *Ifnar2*^−/−^ BM chimeric mice. Graphs show pooled data (mean ± SEM) from two independent experiments. *P < 0.05, **P < 0.01, ***P < 0.001, ns, non-significant, by ANOVA with Kruskal-Wallis test. (D) Experimental schematic of IFNAR blocking during HSV and coinfection. Mice were infected s.c. with HSV, infected i.p. with LCMV on day 2 and injected with anti-IFNAR blocking antibody or isotype control (ISC) antibody on days 2-4. Mice were analyzed on 8 d post-HSV infection. (E-G) Analysis of cellularity of stromal cell subsets (E), HSV-specific CD8^+^ T cells (F) and B cell response (G) in the pLN of HSV and coinfected mice during IFNAR blocking. Graphs show pooled data (mean ± SEM) from 2 independent experiments with 5 mice per group. *P < 0.05, **P < 0.01, ****P < 0.0001, ns, non-significant, by unpaired two-tailed t test (E-G). (H) pLN sections from mice treated with anti-IFNAR blocking antibody or ISC were stained for IgD, CD138, GL7 and Lyve1 and analyzed by confocal microscopy. Data are representative of 2 experiments with 2 mice per group. Scale bar, 200μm.

To better define the impact of IFNAR signals on local LN responses, we administered blocking antibodies against IFNAR (anti-IFNAR) to mice infected with HSV or mice coinfected with LCMV in order to reduce IFNAR signals (Fig. 6D). Anti-IFNAR treatment did not significantly alter responses in mice infected singly with HSV (Fig. 6E). In contrast, anti-IFNAR blockade significantly enhanced the expansion of FRC, LEC, BEC and HEV in coinfected mice (Fig. 6E and Fig. S4F). HSV-specific T cell responses were not altered by anti-IFNAR treatment (Fig. 6F), but T and B cell recruitment, GC B cells and ASC numbers were restored by anti-IFNAR treatment of coinfected mice (Fig. 6G and Fig. S4F). Microscopy revealed that blocking IFNAR signals improved GC formation and CD138^+^ plasma cell numbers (Fig. 6H). Together, these findings show that systemic inflammation resulted in an IFNAR-dependent lymphopenia that reduced cell recruitment to LN draining the site of local infection and impaired LN remodeling and humoral immunity.

## Discussion

Coinfection with multiple pathogens occurs frequently and can alter disease outcomes and subsequent immunity (Mabbott, 2018; Stelekati and Wherry, 2012). Diverse mechanisms likely contribute to altered immunity during pathogen coinfections, yet our understanding of these processes remains incomplete. Here, we show that widespread inflammation triggered by systemic coinfection impairs dLN remodeling and B cell responses during concurrent localised viral infection or immunization. Mechanistically, systemic inflammation results in an IFN-I-dependent lymphopenia and impairs the recruitment of lymphocytes to LN draining sites of peripheral infection, thereby constraining the induction of local humoral immunity.

Recruitment of lymphocytes to inflamed LNs triggers LN expansion in response to infection or immunization (Gregory et al., 2017; Yang et al., 2014). This requires circulating lymphocytes as a source of cells to feed the draining LN. Systemic infections (including malaria and COVID-19) and autoimmune diseases (including systemic lupus erythematosus, SLE) can induce lymphopenia, resulting in the trapping of lymphocytes in secondary lymphoid organs around the body, thereby reducing the circulating pool of lymphocytes. We show that induction of lymphopenia during systemic infection, inflammation or following FTY720 treatment impairs draining LN responses. B cells can coordinate the expansion of the lymphatic network in LNs during inflammation (Angeli et al., 2006; Kumar et al., 2010). We show that removal of B cells impaired the expansion and activation of FRC but had little impact on expansion of LEC or BEC during local HSV infection. The reason for this difference is unclear but may reflect the ability of both CD4^+^ and CD8^+^ T cells to also support LN expansion and remodeling (Gregory et al., 2017).

Our results show that concurrent systemic inflammation impaired lymphocyte recruitment and LN remodeling, which restrained B cell responses but did not impact CTL responses. LN hypertrophy peaks 1-2 weeks after infection, whereas T cells are activated and leave the draining LNs within 1 week (Hor et al., 2015). Conversely, B cells require a prolonged period of activation and maturation within LN in order to form antibody secreting plasma cells and long-lived memory B cells (Cyster and Allen, 2019). This indicates that lymphadenopathy may be critical for the support of B cell responses, probably to support GC reactions and memory B cell formation in the dLN. In support of this, we also observed reduced numbers of T_FH_ cells in coinfected mice. Recruitment of cells to the dLN and expansion of LSC during infection requires IFNAR signaling (Gregory et al., 2017). We did not find a role for direct IFNAR signaling in B cells, which supports a previous study showing that IFNAR^−/−^ B cells did not show increased migration to LN during lymphopenia (Kamphuis et al., 2006). In contrast, extrinsic IFNAR signals influence B cell accumulation in inflamed LNs (Hastey et al., 2014). Whether IFNAR signaling results in the production of other inflammatory mediators (such as IL-6) that further contribute to the sequestration of lymphocytes requires further investigation. Many medically important pathogens cause acute or persistent systemic infections, and evidence suggests that immunity to vaccination or challenge with unrelated pathogens is often impaired during coinfection (Griffiths et al., 2011; Stelekati and Wherry, 2012). Models of viral and bacterial coinfection have revealed increased susceptibility to infection and altered responses. During the early stage of LCMV infection, IFN-I sensitizes mice to bacterial endotoxin (Doughty et al., 2001; Nansen and Randrup Thomsen, 2001), which can lead to NK cell-mediated impairment of the CD8^+^ T cell responses (Straub et al., 2018). Coinfection of mice with two systemic viral pathogens, LCMV and Ectromelia virus (McAfee et al., 2015) or LCMV and Pichinde virus was also found to impair CD8^+^ T cell responses to LCMV (Kenney et al., 2015). In our experiments, LCMV coinfection did not inhibit HSV-specific CD8^+^ T cell responses but rather impaired the induction of local B cell responses. In humans, coinfection may both directly and indirectly affect B cell differentiation and antibody production (Stelekati and Wherry, 2012). Bacterial coinfection of mice with *Streptococcus pneumoniae* was shown to either enhance or inhibit B cell responses to influenza virus depending on the timing of respiratory coinfection (Wu et al., 2015). During the early phase of chronic LCMV infection, virus-specific B cell responses can be inhibited by IFN-I production that results in death of activated B cells and induction of short-lived ASCs at the expense of sustained neutralizing antibody responses (Fallet et al., 2016; Moseman et al., 2016; Sammicheli et al., 2016). Other viruses also directly interfere with B cell responses (Kuka and Iannacone, 2018).

Here, we reveal a novel mechanism by which infection can indirectly interfere with B cell responses. We show that systemic inflammation induced by LCMV infection, the TLR3 agonist Poly(I:C) or the endotoxin LPS induced a lymphopenia that reduced the recruitment of lymphocytes to the dLN during localized infection. This suppression of dLN swelling, stromal cell expansion and B cell immunity suggests that opportunistic localized infections might be harder to eradicate in hosts undergoing systemic inflammatory reactions. Both infectious and non-infectious systemic inflammatory reactions might suppress regional immunity when occurring contemporaneously. In contrast, latent coinfection with murine gammaherpesvirus 68 or murine cytomegalovirus was shown to protect against bacterial infection (Barton et al., 2007), at least transiently (Yager et al., 2009), due to the systemic production of low levels of cytokines. This indicates that the magnitude of systemic inflammation plays important roles in influencing the outcomes of concomitant regional immune responses.

Overall, our data show that systemic inflammation impacts the hosts ability to mount immune responses in LN draining the site of a localized challenge by impairing cell recruitment and restricting tissue remodeling and B cell responses. The prevalence of coinfections in people suggests that it will be important to better define how systemic inflammatory responses in humans impact regional immunity and the progression of diseases that are initiated in barrier tissues.

## Methods

### Mice

C57BL/6, B6.SJL-PtprcaPep3b/BoyJ (CD45.1), OBI x B6.129S7-*Rag1*^*tm1Mom*^/J (OBI.RAG^−/−^)(Dougan et al., 2012), *Ifnar2*^−/−^, B6.129S2-*Ighm*^*tm1Cgn*^/J (μMT^−/−^) and B6.129S4-*Ccr2*^*tm1Ifc*^/J (CCR2^−/−^) mice were bred in the Department of Microbiology and Immunology. Animal experiments were approved by The University of Melbourne Animal Ethics Committee. All mice were sex and aged matched and used between 8-14 weeks old at the beginning of experiments.

### Virus infections

Mice were infected subcutaneously (s.c.) in the footpad with either 5 × 10^4^ plaque-forming units (PFU) of HSV-1 (KOS) or 2 × 10^4^ PFU of LCMV (Armstrong). Mice were infected intranasally with 10^4^ of the recombinant influenza virus X31 expressing the HSV glycoprotein-B-derived epitope gB_498–505_ as described(Davies et al., 2017). For all infections, mice were anaesthetized with isoflurane vaporized in O_2_. Unless stated otherwise mice were coinfected intraperitoneally with 2 × 10^5^ PFU of LCMV Armstrong two days after footpad HSV infection. HSV-1 viral titers were determined in homogenized footpads by PFU assay as described(Jones et al., 2000).

### Stromal cell isolation and flow cytometry

pLN and iLN were harvested and processed as described previously(Gregory et al., 2017). Briefly, LN were teased apart with forceps and incubated at 37°C in RPMI with collagenase D, Dispase and DNase for 25 minutes. Following a second round of digestion for 15 minutes, single cell suspensions were resuspended in FACS buffer (PBS 2% BSA 5mM EDTA) and filtered through 70μM before antibody staining. Antibodies used in this study are listed in Table S1.

Live cells were discriminated with a fixable LIVE/DEAD™ Fixable Near-IR Dead Cell Stain Kit (Thermofischer). Annexin V staining was carried out with Annexin V PE and 7AAD from Biolegend according to the manufacturer’s instructions.

Intracellular staining for KI67 and LCMV were performed using BD Cytofix/Cytoperm™ Fixation/Permeabilization Solution Kit according to the manufacturer’s instructions. Purified LCMV antibody (Clone VL4, BioXcell) was conjugated to Alexa Fluor™ 594 with antibody labelling kit (ThermoFischer) according to the manufacturer’s instructions.

Endogenous HSV and LCMV-specific CD8 T cells were identified with H-2Kb-gB_498-505_ and H-2Db-gp_33-41_ (MBL international)-restricted tetramer staining respectively.

For metabolism analysis, cells were incubated with Mitotracker Deep Red FM, Bodipy FL C 16, CellROX orange reagent or 2-NBDG (all from ThermoFischer) in RPMI supplemented with 10% FCS for 30 minutes at 37°C except for 2-NBDG where glucose-free RPMI (ThermoFischer) was used instead.

Cells were enumerated by adding SPHERO calibration particles (BD Biosciences) to each sample before acquisition using FACSCantoII or FACSFortessa (both BD) and Flowjo software was used for analysis.

### Immunomodulatory treatments

For CD8^+^ T cell depletion, mice were injected i.p. twice with 100 μg of anti-CD8 antibody (clone 2.43), 24 hours apart. B cells were depleted with a single injection of 50 μg of anti-mouse CD20 antibody (clone 5D2, Genentech) prior HSV infection. To block lymphocyte egress, FTY720 (2-amino-2-(2-[4-octylphenyl]ethyl)-1,3-propanediol; Cayman chemical, #402616-26-6) was dissolved in 2% cyclodextrin (Sigma) in PBS and i.p. administered daily in mice at a dose of 1mg/kg from day 1 to day 7 post-HSV infection. Control mice were administered with 2% cyclodextrin only. To analyze the effect of systemic inflammation on local immune responses, HSV-infected mice were injected i.p. twice with either 50 μg Polyinosinic:polycytidylic acid high molecular weight (Poly(:IC), InvivoGen) or 5 μg of LPS (Sigma), 24 hours apart. IFNAR in vivo blocking during coinfection was performed by injecting infected mice with 500 μg of anti-IFNAR-1 (clone MAR1-5A3, from BioXcell) or isotype control antibody (clone MOPC-21, BioXcell) for two consecutive days and 250 μg on the third day. Immunization of mice was carried out by injecting 100μg of OVA (Sigma) emulsified in CFA (Sigma) s.c.

### Lymphocyte recruitment analysis

C57BL/6-CD45.2 mice were infected s.c. with HSV and two days later injected i.v. with a mixture of 8 million total lymphocytes from pooled LN and spleen from C57BL/6-CD45.1 mice immediately prior to coinfection with LCMV. HSV draining and non-draining LN were analyzed 2 days post LCMV infection.

### Generation of bone marrow chimeric mice

To generate IFNAR deficient B cell mice, C57BL/6-Ly5.1 recipient mice were lethally irradiated with two doses of 550 rads, 3 h apart, followed by reconstitution with a 80/20 mixture of 8 × 10^6^ μMT^−/−^ / WT or μMT^−/−^ / *Ifnar2*^−/−^ donor bone marrow cells and maintained in antibiotic water (neomycin and polymyxin B) for 4 weeks. To analyze the contribution of IFNAR signalling on hematopoietic and non-hematopoietic compartments, C57BL/6-Ly5.2 WT and *Ifnar2*−/− mice were irradiated and reconstituted with either 8 × 106 BM cells from WT or *Ifnar2*−/− donor mice. All recipients were used at least 8 weeks post-reconstitution.

### Immunofluorescence and confocal microscopy

Lymph nodes were harvested and fixed in periodate-lysine-paraformaldehyde (PLP) fixative for 4 hours, incubated in 30% sucrose and embedded in OCT freezing media. LN were harvested and immediately embedded in OCT freezing media. Tissue sections were cut at 16 μm thickness with a cryostat (Leica CM3050S) and air-dried before being fixed in acetone for 5 min, dried, and then blocked for 30 min (Protein Block X0909, DAKO) at room temperature (RT). Sections were staining with primary antibodies for 1 hour at RT, washed in PBS and when required stained with secondary antibodies for 45 minutes at RT. Stained sections were mounted in ProLong Gold antifade reagent (Invitrogen), and acquired on a LSM780 or LSM710 confocal microscope (Carl Zeiss) and images processed with Imaris (Bitplane). Antibodies used for histology are listed in Table S1.

### Detection of HSV-1 IgG by ELISA

Nunc MaxiSorp round-bottom 96-well ELISA microtiter plates (Thermo Scientific) were coated overnight at 4°C with 10 μg/ml HSV-1 inactivated Vero cell extract (Advanced Biotechnologies). Unbound protein was washed away (PBS 0.05% Tween20) and serially diluted serum samples (PBS 0.5% BSA) were plated and incubated at RT for two hours. Bound mouse IgG Abs were detected using donkey anti-mouse IgG-HRP and visualized using o-Phenylenediamine. The OD readings were determined at 450/492 nm. Endpoint titers of anti-HSV-1 were calculated by using cut-off values defined as double the OD of non-infected serum control.

### Statistical analysis

Graphs and statistics were generated using Prism 8 (GraphPad). Samples were tested for normality and two groups were compared using two-tailed Mann-Whitney U-test or unpaired t-test. Multiple groups were analyzed with one-way Anova followed by Tukey’s post-test comparison or Kruskal-Wallis, based on Gaussian distribution. All graphs depict means ± SEM.

**Table S1.**
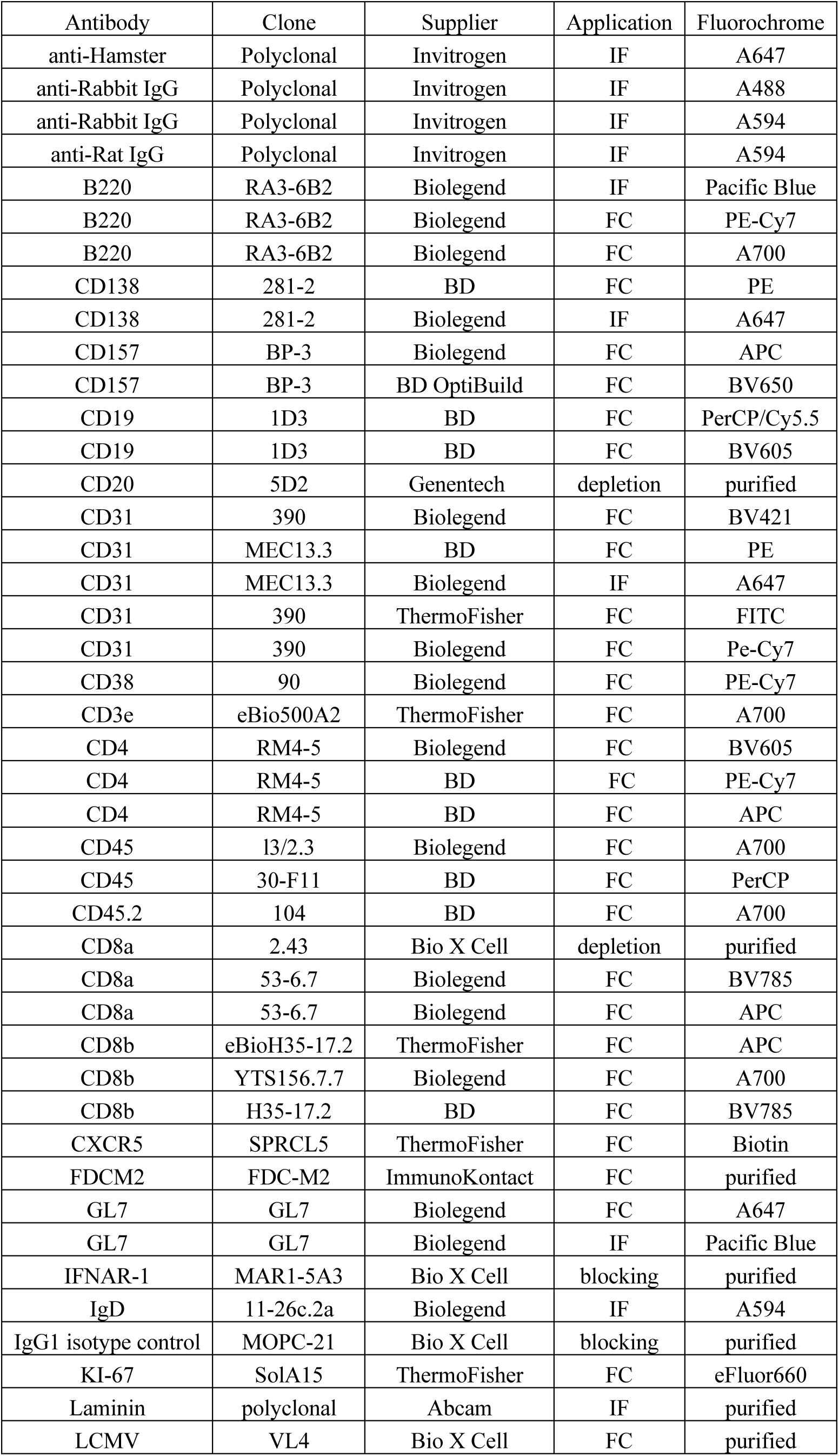
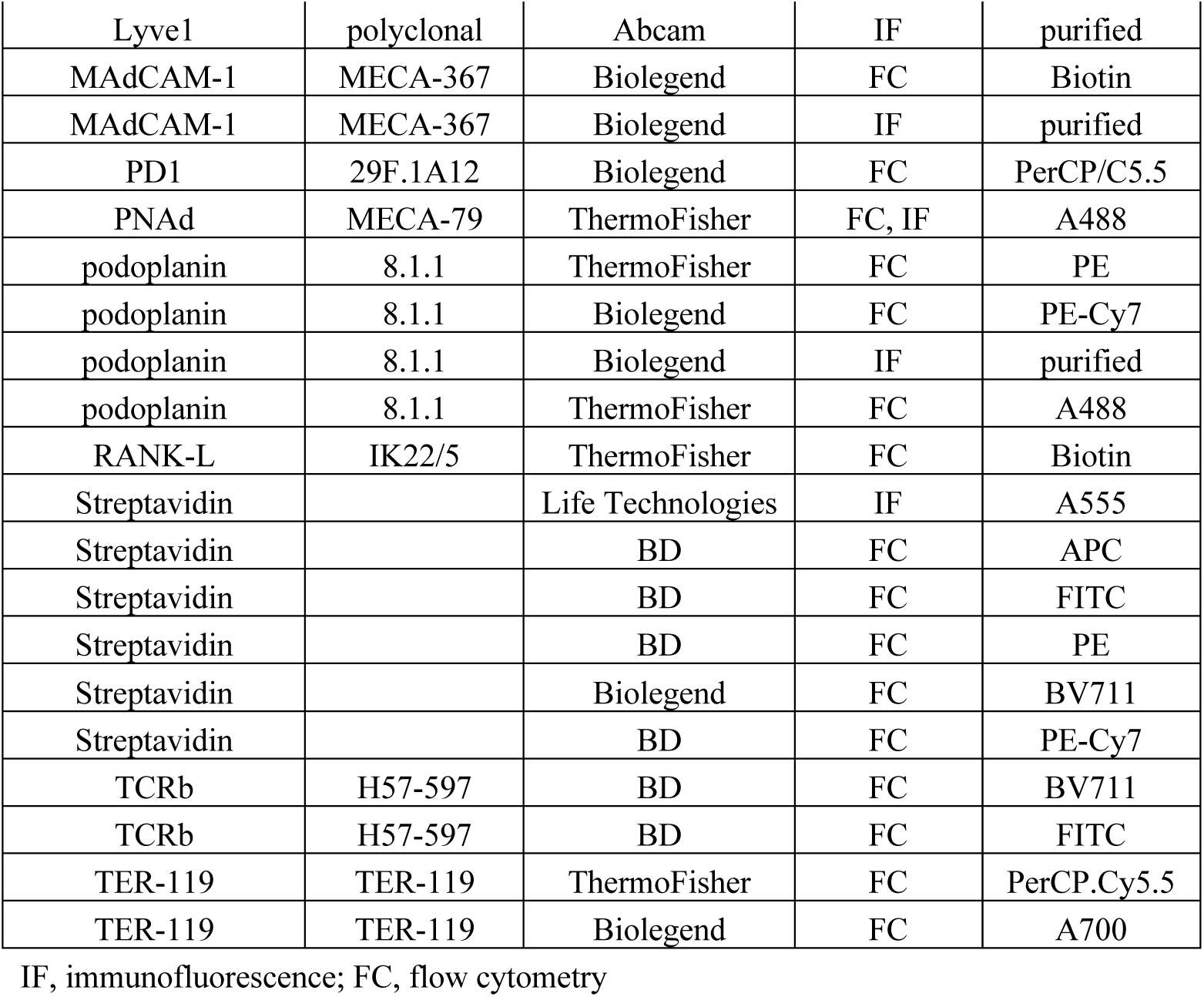
Antibodies used for immunofluorescence staining, flow cytometry, blocking and depletion experiments.

| Antibody | Clone | Supplier | Application | Fluorochrome |
| --- | --- | --- | --- | --- |
| anti-Hamster | Polyclonal | Invitrogen | IF | A647 |
| anti-Rabbit IgG | Polyclonal | Invitrogen | IF | A488 |
| anti-Rabbit IgG | Polyclonal | Invitrogen | IF | A594 |
| anti-Rat IgG | Polyclonal | Invitrogen | IF | A594 |
| B220 | RA3-6B2 | Biolegend | IF | Pacific Blue |
| B220 | RA3-6B2 | Biolegend | FC | PE-Cy7 |
| B220 | RA3-6B2 | Biolegend | FC | A700 |
| CD138 | 281-2 | BD | FC | PE |
| CD138 | 281-2 | Biolegend | IF | A647 |
| CD157 | BP-3 | Biolegend | FC | APC |
| CD157 | BP-3 | BD OptiBuild | FC | BV650 |
| CD19 | 1D3 | BD | FC | PerCP/Cy5.5 |
| CD19 | 1D3 | BD | FC | BV605 |
| CD20 | 5D2 | Genentech | depletion | purified |
| CD31 | 390 | Biolegend | FC | BV421 |
| CD31 | MEC13.3 | BD | FC | PE |
| CD31 | MEC13.3 | Biolegend | IF | A647 |
| CD31 | 390 | ThermoFisher | FC | FITC |
| CD31 | 390 | Biolegend | FC | Pe-Cy7 |
| CD38 | 90 | Biolegend | FC | PE-Cy7 |
| CD3e | eBio500A2 | ThermoFisher | FC | A700 |
| CD4 | RM4-5 | Biolegend | FC | BV605 |
| CD4 | RM4-5 | BD | FC | PE-Cy7 |
| CD4 | RM4-5 | BD | FC | APC |
| CD45 | 13/2.3 | Biolegend | FC | A700 |
| CD45 | 30-F11 | BD | FC | PerCP |
| CD45.2 | 104 | BD | FC | A700 |
| CD8a | 2.43 | Bio X Cell | depletion | purified |
| CD8a | 53-6.7 | Biolegend | FC | BV785 |
| CD8a | 53-6.7 | Biolegend | FC | APC |
| CD8b | eBioH35-17.2 | ThermoFisher | FC | APC |
| CD8b | YTS156.7.7 | Biolegend | FC | A700 |
| CD8b | H35-17.2 | BD | FC | BV785 |
| CXCR5 | SPRCL5 | ThermoFisher | FC | Biotin |
| FDCM2 | FDC-M2 | ImmunoKontakt | FC | purified |
| GL7 | GL7 | Biolegend | FC | A647 |
| GL7 | GL7 | Biolegend | IF | Pacific Blue |
| IFNAR-1 | MAR1-5A3 | Bio X Cell | blocking | purified |
| IgD | 11-26c.2a | Biolegend | IF | A594 |
| IgG1 isotype control | MOPC-21 | Bio X Cell | blocking | purified |
| KI-67 | SolA15 | ThermoFisher | FC | eFluor660 |
| Laminin | polyclonal | Abcam | IF | purified |
| LCMV | VL4 | Bio X Cell | FC | purified |

| Lyve1 | polyclonal | Abcam | IF | purified |
| --- | --- | --- | --- | --- |
| MAdCAM-1 | MECA-367 | Biolegend | FC | Biotin |
| MAdCAM-1 | MECA-367 | Biolegend | IF | purified |
| PD1 | 29F.1A12 | Biolegend | FC | PerCP/C5.5 |
| PNAd | MECA-79 | ThermoFisher | FC, IF | A488 |
| podoplanin | 8.1.1 | ThermoFisher | FC | PE |
| podoplanin | 8.1.1 | Biolegend | FC | PE-Cy7 |
| podoplanin | 8.1.1 | Biolegend | IF | purified |
| podoplanin | 8.1.1 | ThermoFisher | FC | A488 |
| RANK-L | IK22/5 | ThermoFisher | FC | Biotin |
| Streptavidin |  | Life Technologies | IF | A555 |
| Streptavidin |  | BD | FC | APC |
| Streptavidin |  | BD | FC | FITC |
| Streptavidin |  | BD | FC | PE |
| Streptavidin |  | Biolegend | FC | BV711 |
| Streptavidin |  | BD | FC | PE-Cy7 |
| TCRb | H57-597 | BD | FC | BV711 |
| TCRb | H57-597 | BD | FC | FITC |
| TER-119 | TER-119 | ThermoFisher | FC | PerCP.Cy5.5 |
| TER-119 | TER-119 | Biolegend | FC | A700 |

IF, immunofluorescence; FC, flow cytometry

## Acknowledgments

We thank Paul Hertzog (Hudson Institute, Australia) for anti-IFNAR antibody, Hidde Ploegh (Whitehead Institute, Cambridge, MA) for OBI mice, Genentech for anti-mouse CD20 antibody, the Biological Optical Microscopy Platform (BOMP) for support, and Yu Kato for advice on HSV ELISA.

## Author contributions

Conceptualization, YOA and SNM; Methodology, YOA and SD; Investigation, YOA, SD, and SLP; Writing, YOA and SNM; Resources, PJH, LKM and WRH; Visualization, YOA and SNM; Funding acquisition, SNM.

## Supplemental Figures

**Fig. S1.**
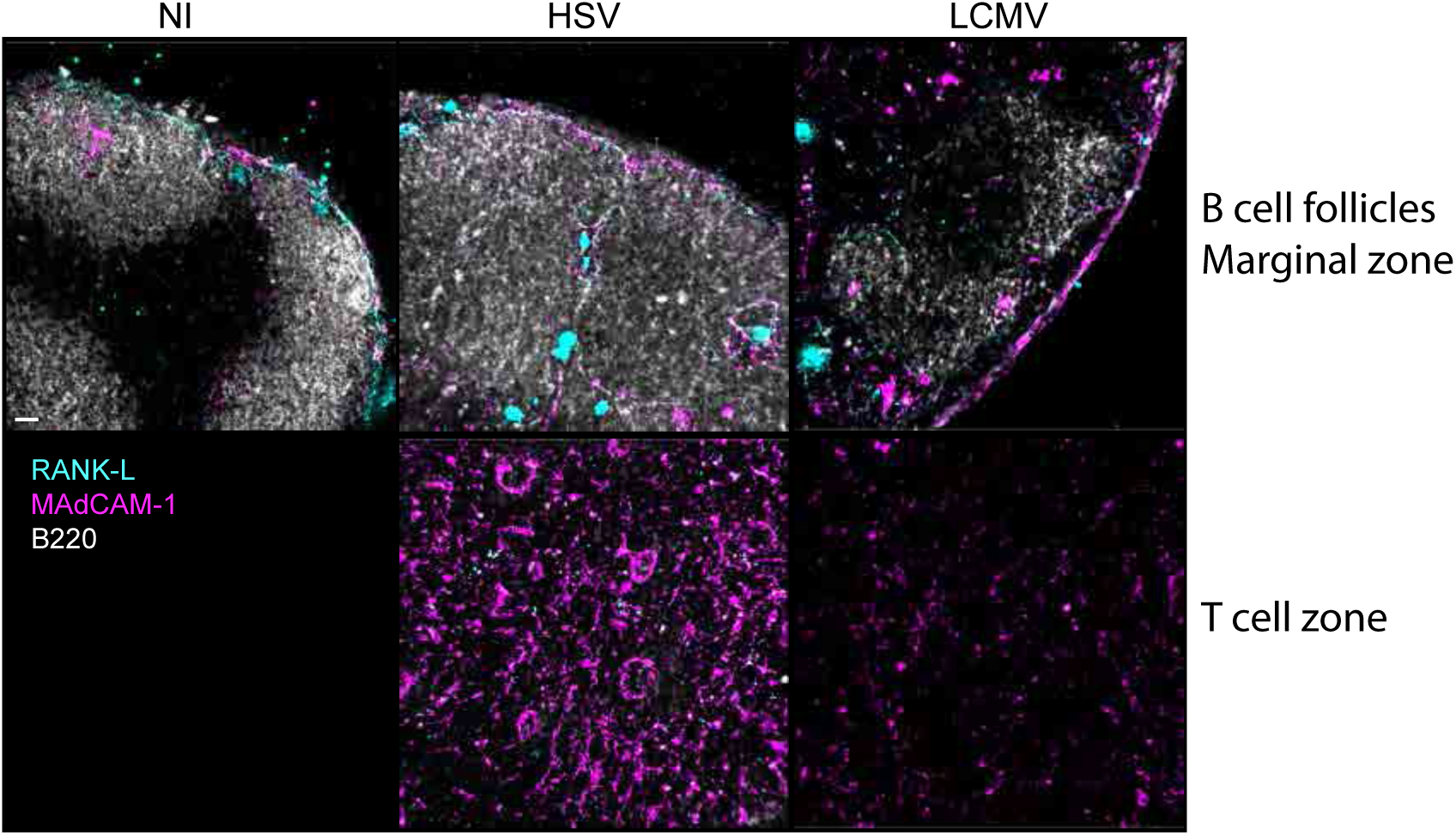
FRC upregulate MAdCAM-1 expression during HSV infection. pLN sections from NI, HSV or LCMV infected mice were stained for B220, RANK-L and MAdCAM-1, and analyzed by confocal microscopy. Data are representative of 2 experiments with 3 mice per group. Scale bar, 40μm.

**Fig. S2.**
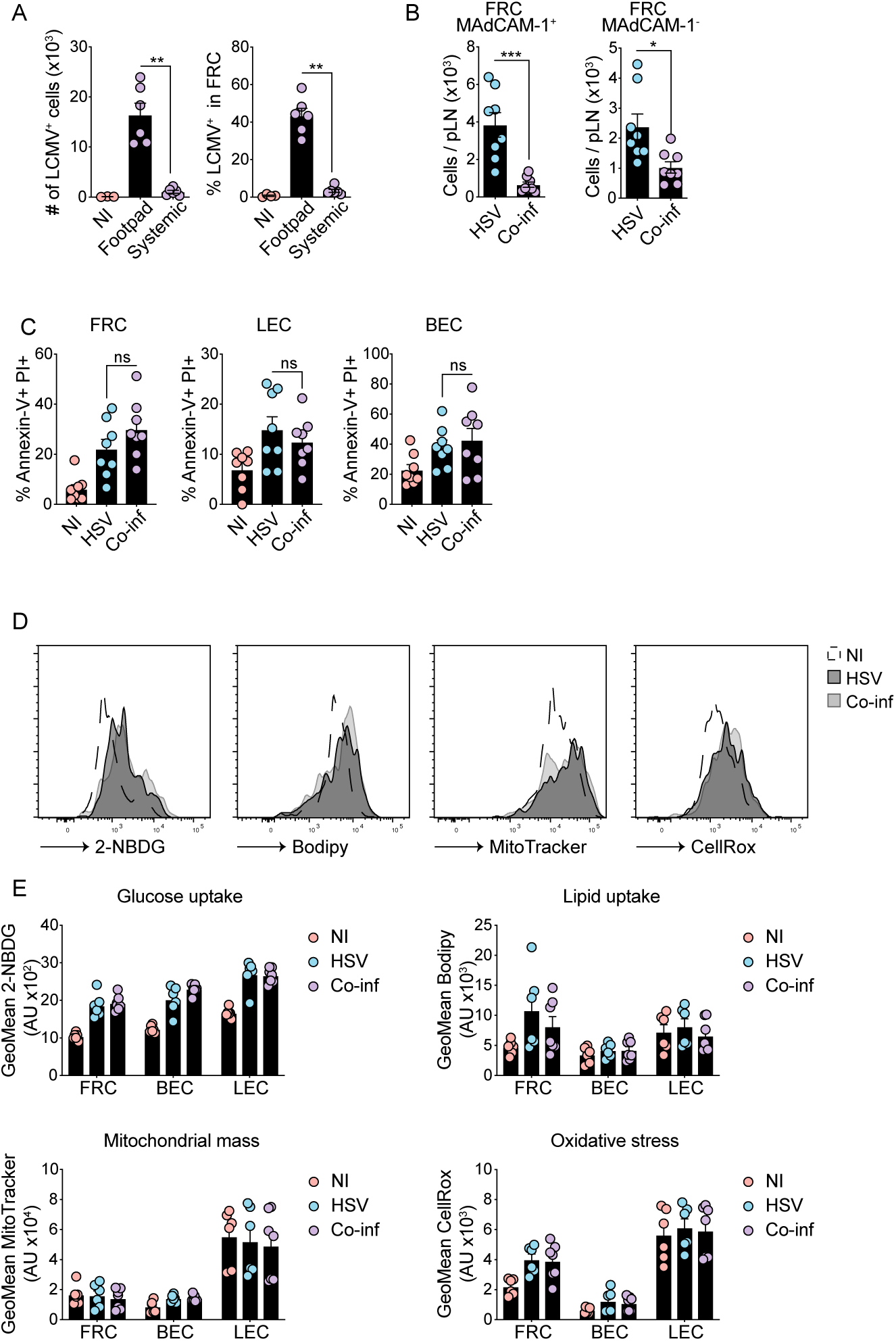
Systemic coinfection impairs stromal cell expansion. (A) Mice were infected with LCMV s.c. or i.p. and pLN harvested 3 d post-infection. LCMV was detected by intracellular staining and analyzed by flow cytometry. Left graph displays total LCMV infected cells, right graph the percentage of infected FRC. Graphs show pooled data (mean ± SEM) from 2 independent experiments with 3 mice per group. **P < 0.01, by Mann-Whitney test (B) Mice were infected with HSV or coinfected as described in Fig. 3. Numbers of FRC expressing or not MAdCAM-1 in pLN were analyzed by flow cytometry. Graphs show pooled data (mean ± SEM) from 2 independent experiments with 4 mice per group. *P < 0.05, *** P < 0.001, by unpaired two-tailed t test. (C) pLN from HSV and coinfected mice were harvested 5 d post-infection and analyzed for Annexin V and PI expression on stromal cells by flow cytometry. iLN from naïve mice were included as controls. Graphs show pooled data (mean ± SEM) from 2 independent experiments with 4 mice per group. ns, non-significant, by by unpaired two-tailed t test (D-E) Metabolism of stromal cells during infection. Mice were infected as in Fig. 3 and pLN harvested 8 days post-infection. iLN from naïve mice were used as controls. Cells were incubated with the fluorescent glucose analog 2-NBDG, the fluorescent fatty acid analog BODIPY FL C16, Mitotracker or CellRox before analysis by flow cytometry. Representative histograms on FRC (D) and analysis of geometric mean values on stromal cell subsets (E). Graphs show pooled data (mean ± SEM) from two independent experiments with 3-4 mice per group.

**Fig. S3.**
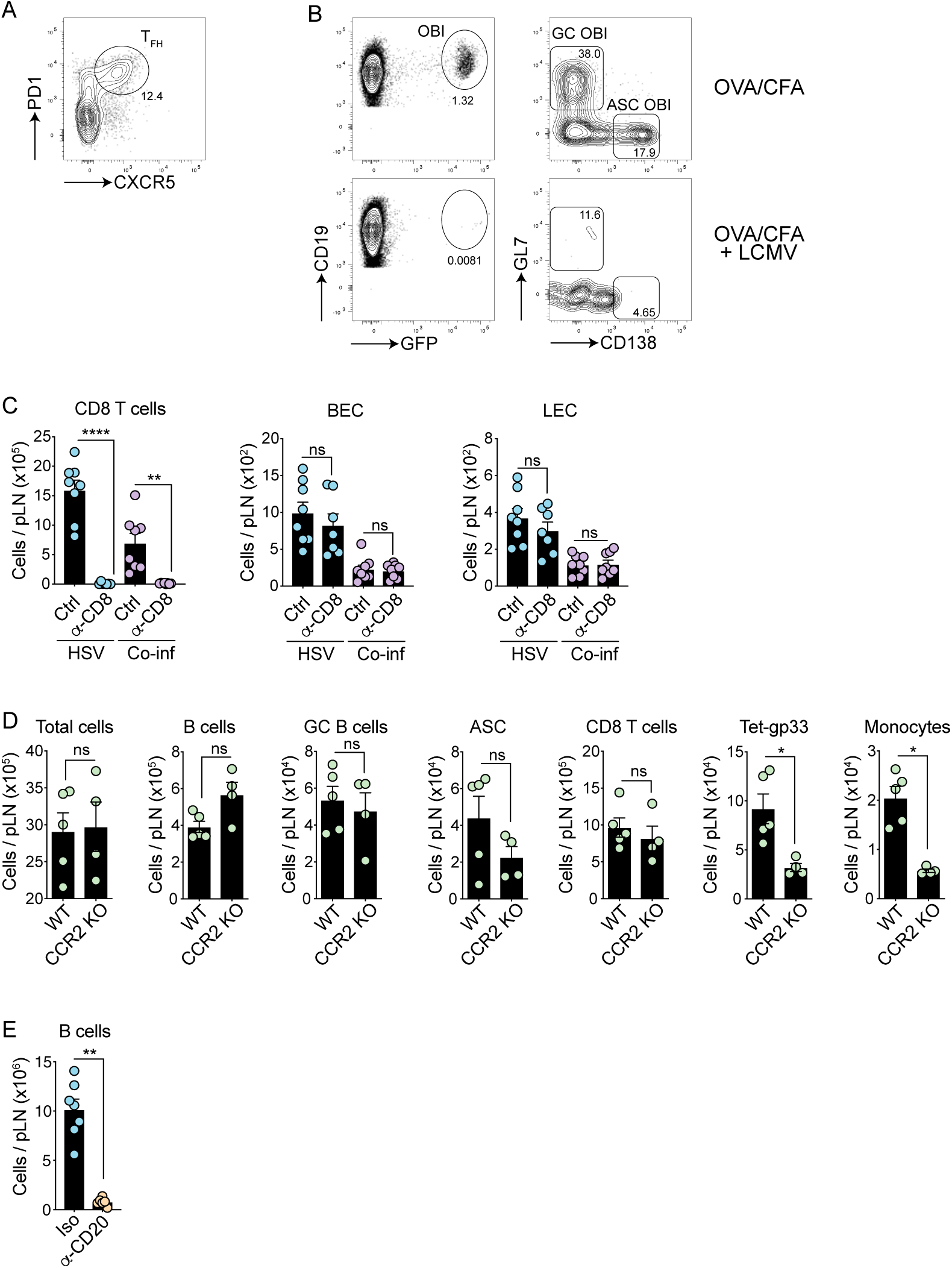
Lymphopenia suppresses LN expansion and B cell responses. (A) Gating strategy to identify T_FH_ in infected mice. CD4^+^ T cells were analyzed for CXCR5 and PD1 expression and TFH cells defined as CD4^+^ CXCR5^+^ PD1^+^ cells. (B) OBI B cell response in the pLN. Mice were injected i.v. with transgenic OBI-GFP cells and two days later injected s.c. with an emulsion of OVA/CFA. Mice were infected i.p. with LCMV 2 d post immunisation and pLN harvested 8 d post OVA/CFA injection. Gating strategy to identify OBI cells in the pLN. CD19^+^ CD3ε^−^ B cells were analyzed for GFP expression and OBI cells analyzed for GL7 and CD138 expression to identify GC OBI and ASC OBI. (C) Experimental procedure as in Fig. 5A. Absolute numbers of CD8^+^ T cells BEC and LEC in the pLN of mice infected with HSV or coinfected with or without CD8^+^ T cells. Graphs show pooled data (mean ± SEM) from 2 independent experiments with 3-4 mice per group. **P < 0.01, ****P < 0.0001, ns, non-significant, by ANOVA with Krukal-Wallis test (D) Monocytes are not required for CD8^+^ and B cell responses during LCMV infection. WT and CCR2 KO mice were infected with LCMV s.c. and pLN harvested 8 d post-infection. B cell and T cell responses were analyzed by flow cytometry. Graphs show data from 1 representative experiment out of 2 (mean ± SEM) with 4-5 mice per group. (E) B cell depletion following CD20 antibody depleting antibody injection. See Fig. 5F for experimental procedure. *P < 0.05, **P < 0.01, ns, non-significant, by Mann-Whitney test.

**Fig. S4.**
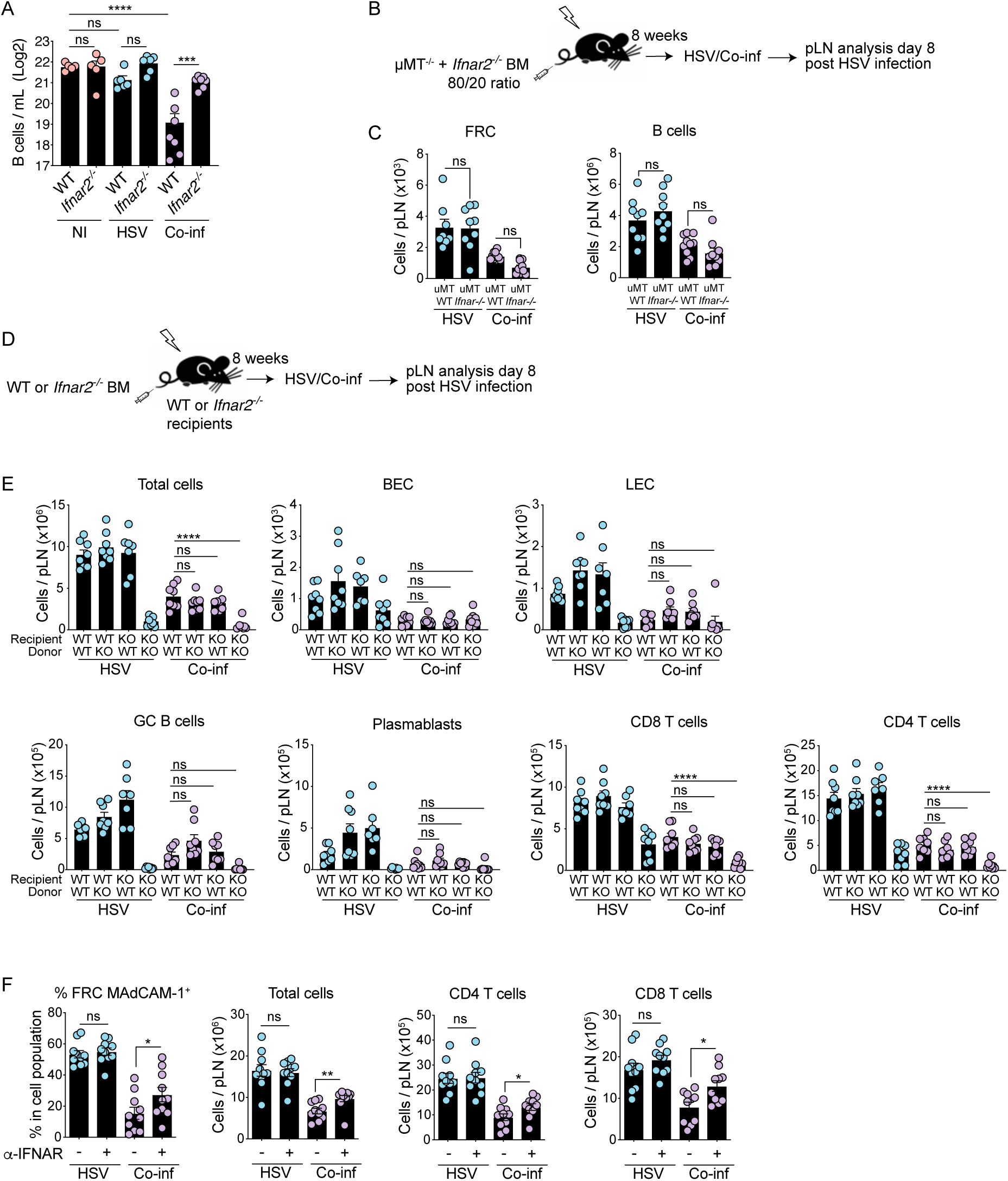
IFN-I induces local LN suppression via B cell-extrinsic IFNAR signals. (A) B cell cellularity in the blood of infected mice. WT or *Ifnar2*^*−/−*^ mice were infected with HSV s.c. and the following day infected with LCMV i.p. Blood was collected on day 3 and B cell numbers analyzed by flow cytometry. Graph shows pooled data (mean ± SEM) from 2 independent experiments with 2-4 mice per group. **P < 0.01, *** P < 0.001, ****P < 0.0001, ns, non-significant, by ANOVA with Dunnett’s multiple comparisons test. (B) Experimental schematic of generating IFNAR deficient B cells. Lethally irradiated mice received a mixture of 80/20 bone marrow cells from μMT + *Ifnar2* or μMT + WT donor mice. 8 weeks post reconstitution, mice were HSV and coinfected as described in Fig. 3A. (C) Absolute numbers of FRC and B cells in the pLN of HSV and coinfected mice in IFNAR B cell deficient mice. Graphs show pooled data (mean ± SEM) from two independent experiments with 4-5 mice per group. *P < 0.05, **P < 0.01, ***P < 0.001, ns, non-significant, by ANOVA with Kruskal-Wallis test. (D) Experimental schematic of generating IFNAR BM chimeras. Lethally irradiated WT or *Ifnar2*^*−/−*^ recipient mice received BM cells from WT or *Ifnar2*^*−/−*^ donor mice. 8 weeks post reconstitution, mice were HSV and coinfected. (E) Cell numbers in the pLN of HSV and coinfected mice. Graphs show pooled data (mean ± SEM) from two independent experiments. *P < 0.05, **P < 0.01, ***P < 0.001, ns, non-significant, by ANOVA with Kruskal-Wallis test. (F) Mice were infected as in Fig. 6D. Analysis of cellularity of total cells, HEV, CD4^+^ and CD8^+^ T cells in the pLN of HSV and coinfected mice during IFNAR blocking. Graphs show pooled data (mean ± SEM) from 2 independent experiments with 5 mice per group. *P < 0.05, **P < 0.01, ***P < 0.001, ns, non-significant, by unpaired t test.

